# Crystal Structure of 3-Hydroxypropionyl-CoA Synthetase (ADP-forming) from *Nitrosopumilus maritimus*

**DOI:** 10.1101/2025.09.22.677778

**Authors:** Jerome Johnson, Bilge Tosun, Merve Yilmaz, Bradley B. Tolar, Yasuo Yoshikuni, Christopher A. Francis, Tzanko Doukov, Shun Yokoi, Soichi Wakatsuki, Hasan DeMirci

## Abstract

The 3-hydroxypropionate/4-hydroxybutyrate (3HP/4HB) cycle in thaumarchaeota contributes significantly to global organic carbon fixation as the most energetically efficient aerobic carbon fixation pathway. The thaumarchaeal 3-Hydroxypropionyl-CoA Synthetase (ADP-forming; Nmar_1309) is crucial to this efficiency, utilizing ATP to ADP catalysis. This first reported structure of Nmar_1309 reveals a homodimer with a unique subdomain organization ([3-4-1-2-5]) and a distinct linker between subdomains 4 and 1. The presence of bound substrates including 3-hydroxypropionate, non-hydrolyzable ATP (ADPNP), and a phosphate suggests an intermediate state mimicking a reaction step immediately preceding the formation of a 3-hydroxypropionyl-phosphohistidine. Conformational differences were observed between the two chains of the homodimer, likely influenced by the binding of a single ADPNP molecule in one chain. Phylogenetic analysis suggests that while 4HB synthetases may have evolved earlier in the evolutionary timeline, 3HP synthetases in Thaumarchaeota may have occurred after the Great Oxygenation Event. These structural data provide further characterization of the 3HP/4HB cycle and, in conjunction with the structure of 4-hydroxybutyryl-CoA synthetase, Nmar_0206, provide baseline structures of the key ADP-forming Acyl-CoA synthetases within this pathway.

## Introduction

The looming climate crisis has been accelerated with reports that between January 2023 and January 2024 the average surface temperature of the earth surpassed a critical threshold of 1.5°C^1^. To combat increased greenhouse gasses, the study of biological processes and enzymes uniquely suited to fix atmospheric carbon has become an international endeavor. One such system, the Thaumarchaeotal 3-hydroxypropionate/4-hydroxybutyrate cycle (3HP/4HB cycle), has been identified as the most efficient aerobic carbon fixation cycle currently described^2^. This cycle, which fixes two carbon dioxide molecules in the form of bicarbonate and leads to the formation of the major end products acetyl- and succinyl-CoA, passes through the important intermediates 3-hydroxypropionate (3HP) and 4-hydroxybutyrate (4HB) (see Fig. 1). Significant variations have been found between Thaumarchaeotal and Crenarchaeotal versions of these enzymes which result in stark differences in energetic efficiency^2,3,4,5,6^. Structural characterization of these enzymes will inform engineering efforts to develop improvements on natural sequestration methods.

**Fig. 1:**
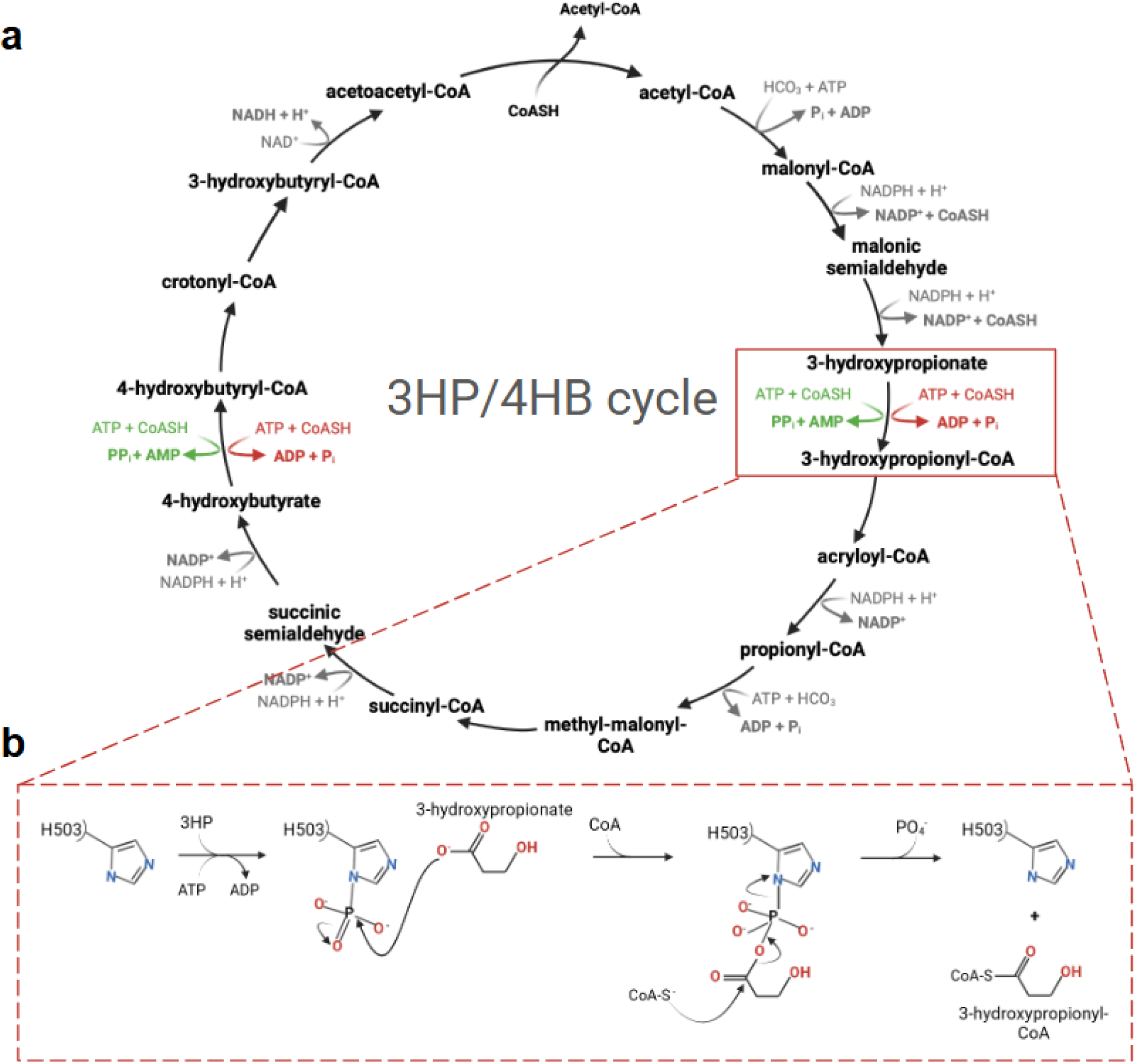
The reactions involved in the HP/HB cycle for crenarchaeal and thaumarchaeal variants. **a** The steps influencing the energy efficiency of the crenarchaeal 3HP/4HB cycle, specifically for *M. sedula*, are highlighted in green while those for thaumarchaeota and *N. maritimus* variant are indicated in red. **b** Simplified reaction mechanism of Nmar_1309.

Within the oligotrophic open oceans, Thaumarchaeota may fix as much as 1% of all the biologically fixed carbon^7^. The (ADP-forming) 3-hydroxypropionyl-CoA synthetase (Nmar_1309, from the representative archaeon *Nitrosopumilus maritimus*) is a critical enzyme in their success as alternatives to the cycle utilize a less energy efficient, AMP-forming version of the enzyme. Nmar_1309 is a member of the (ADP-forming) Acyl-CoA Synthetase superfamily (ACD) of enzymes, each member of which catalyzes the formation of an acyl-CoA and ADP using ATP, CoA and an acyl containing compound as substrate^8^. Nmar_1309 and Nmar_0206 (4-hydroxybutyrl-CoA synthetase) are two of a number of ACDs identified within the *N. maritimus* genome. Although capable of catalyzing multiple substrates, their low catalytic efficiency relative to 3HP supported the identification of Nmar_1309 as a 3-hydroxypropionyl-CoA synthetase.

Succinyl-CoA synthetase (SCS) may be seen as a model for the ACD superfamily, as characterization of the ACD reaction mechanism and structural characteristics for ACDs were first identified in the SCS enzyme^9^. ACD structures consist of CoA-binding subdomains 1 and 2; ATP-binding domains 3 and 4; and CoA-ligase subdomain 5. In short, the reaction mechanism involves the dephosphorylation of the gamma phosphate of a bound ATP molecule resulting in the phosphate covalently bonded and passed to 1 or 2 histidine residues^10^. This was proposed as members of the super family contain ATP-binding sites at varying distances from the active site. Structures with a relatively larger distance between the ATP-binding and active site, such as 31.5 Å within the larger 4-hydroxypropionyl-CoA synthetase compared to ∼20 Å within the smaller SCS, require an additional histidine within the ATP-binding site to support catalysis. Regardless, due to the distances between the ATP binding site and the active site, a ‘swinging-loop’ containing the primary histidine bridges the distance between the two sites. After phosphorylation within the ATP binding site, this ‘swinging-loop’ returns to the active site to bind the acyl-group of the substrate molecule. This forms a acyl-phosphohistidine from which CoA may perform a nucleophilic attack with phosphate acting as a leaving group during the formation of the product Acyl-CoA (see Fig. 1)^11^.

This requirement to bridge large distances with flexible loops is a consequence of the remarkable structural diversity within the ACDs, which often arises from subdomain shuffling. This shuffling results not only in changes to the order of the protein primary structure (amino acid sequence) but also in oligomerization state, linkage and surface interaction variations^3^. The domain organization of ACD superfamily proteins can be described following nomenclature standardized by SCS: [alpha(1,2)/beta(3,4,5)] for SCS^11^, [alpha(1,2,5)/beta(3,4)] for Acetyl-CoA synthetase (pdb: 4XYM)^12^, [1,2,5,3,4] for 4-hydroxybutyryl-CoA synthetase (Nmar_0206 from the same cycle)^3^ and [3,4,1,2,5] for Nmar_1309. From this organization, ACDs oligomerize into heterodimeric, homodimeric and heterotetrameric structures. These are found throughout all domains of life. To address how and why this structural diversity evolved, this study reveals the molecular architecture of Nmar_1309, providing the first structural snapshot of this unique domain arrangement within the ACD superfamily.

## Materials and Methods

### Cloning

The *Nitrosopumilus maritimus* Nmar_1309 (3-hydroxypropionyl-CoA synthetase) gene sequence with an N-terminal hexahistidine tag was designed and codon-optimized using Genscript BioTech trademark software before synthesis for Ni-NTA affinity purification. Nmar_1309 was then cloned into the pET28a vector using *Nde*I and *Bam*HI endonuclease restriction sites, and transformed into *E. coli* strain BL21(Rosetta-2). Transformed cells were selected for by growth in LB agar plates containing kanamycin (50 µg/ml) and chloramphenicol (50 µg/ml) at 37℃.

### Protein Expression and Purification

Expression of Nmar_1309 was performed using the BL21(Rosetta-2) *E. coli* transformed cells. Cultures were grown overnight in LB media and then diluted 1:100 into 2 L cultures. The cell cultures were grown to an optical density of 0.8 at 600 nm and then expressed overnight at 18°C following induction with 0.4 mM IPTG. Cell pellet was then obtained following centrifugation at 3700 rpm, resuspended in a lysis buffer (pH 8.5 50 mM Tris, 300 mM NaCl, 5% v/v glycerol supplemented with 0.01% Triton X-100), and sonicated. Soluble protein was maintained at 4°C, purified using Ni-NTA affinity resin (GE Healthcare) and concentrated to 10 mg/ml using 50 kDa Amicon concentrators. The column was equilibrated with pH 8.5 HisA (containing 300 mM NaCl and 20 mM Tris). A column wash was performed using pH 8.5 HisA and eluted with pH 8.5 HisB (containing 500 mM imidazole, 300 mM NaCl, 50 mM Tris). Following purification, the His_6_-tag was removed overnight at 4°C by thrombin digestion in the presence of 5 mM beta-mercaptoethanol. Reverse Ni-NTA was then performed to remove the thrombin-cleaved His_6_-tag . Nmar_1309 purity was confirmed through SDS-PAGE. Following purification, 20 mL of 50% glycerol was added to 30 mL of the enzyme elutant for long-term storage.

### Crystallization

Crystallization screens were performed using 72-well Terasaki microbatch plates. 0.83 µL of purified Nmar_1309 were pipetted into the bottom of the sitting drop and mixed with an additional 0.83 µL of ∼3500 commercially available sparse-matrix crystallization screening conditions^13^, followed by a 16.6 µL application of Paraffin oil. Plates were stored in styrofoam containers at room temperature and crystals of Nmar_1309 were obtained 1-2 weeks after initial crystallization screenings. Protein crystals were obtained in a solution of 2.5 µL of 200 mM Magnesium chloride hexahydrate, 100 mM Tris pH 8.5, 7% (v/v) PEG 6000 with 2.5 µL of the enzyme solution, covered with ∼50 µL of Paraffin oil.

### Data Collection and Processing

Protein crystals of Nmar_1309 in 20% v/v glycerol were prepared for crystallography by flash freezing in liquid nitrogen. The data was collected at the Advanced Light Source Beamline 5.0.2 at the Lawrence Berkeley National Laboratory, Berkeley, California, USA. The detector distance for the enzyme structure was set at 400 mm with an exposure time of 0.2 s per frame, with the X-ray energy set to 12.65 keV. The diffraction data were collected to 2.8 Å resolution at 100 K. The crystal belongs to the space group P2_1_2_1_2_1_ with unit cell dimensions a = 81.88 Å, b = 125.59 Å, c =137.76 Å, α = 90, β = 90, and γ =90 [see Table 1]. The diffraction data were processed with the XDS package^14^ for indexing and scaled by using XSCALE. STARANISO was used to further improve the data^15^.

**Table 1.**
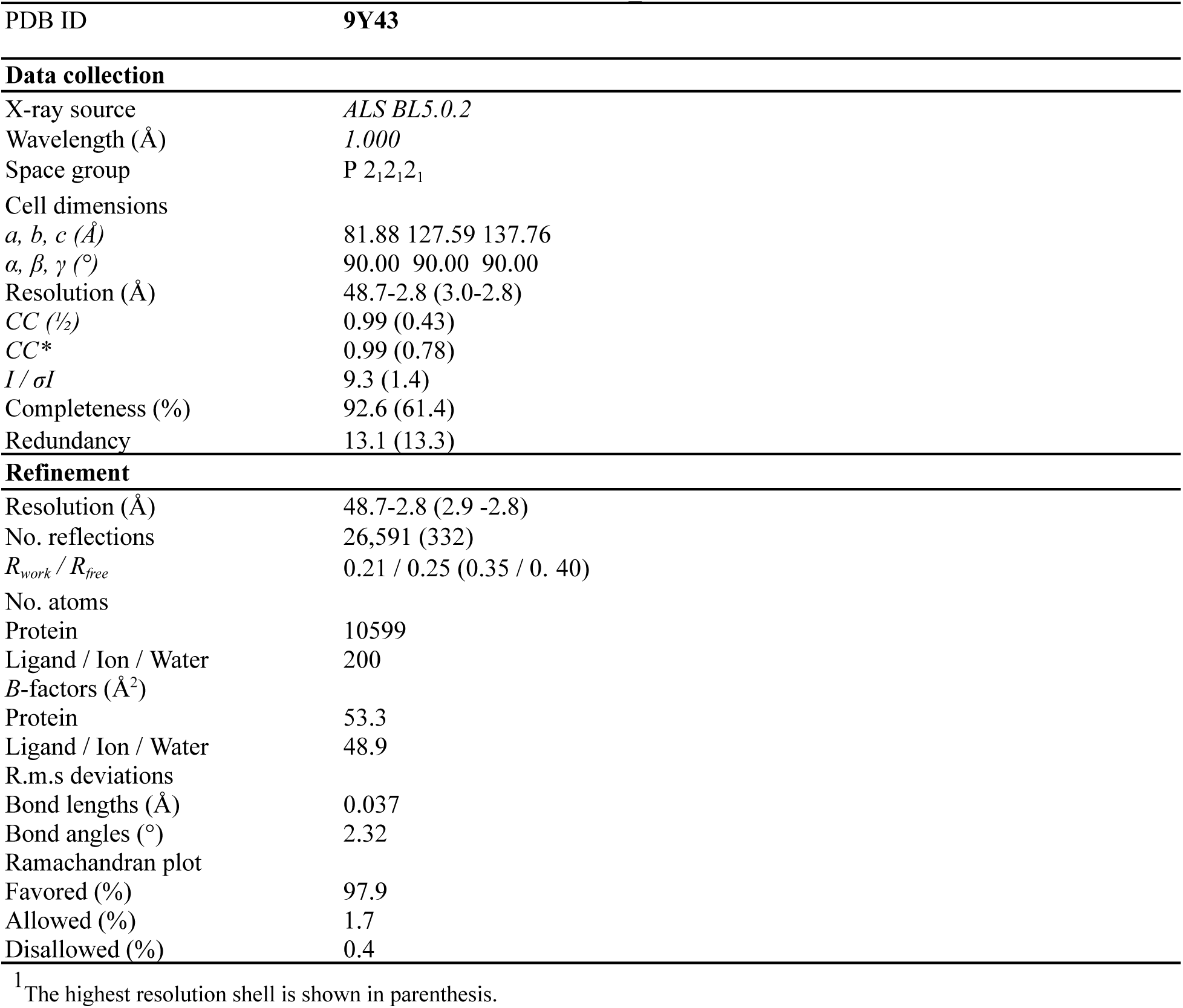
Data collection and refinement statistics for Nmar_1309.

### Structure Determination and Refinement

An AlphaFold2 generated model was used as a search model for automated molecular replacement using *PHASER* within the Phenix software suite^16^. This was followed by simulated-annealing, individual coordinate and TLS parameters refinement. Coot^17^ (version 0.9.8.1) was then used to confirm residue and water positions, and addition of residues missing in the electron density map. Further refinement was performed to 2.8 Å resolution within Coot while maintaining positions with strong difference densities and Ramachandran statistics for the structure were optimized to 97.9/1.6/0.4% (most favored/additionally allowed/disallowed) [see Table 1 for full refinement statistics]. The final R-work and R-free were 0.21 and 0.25, respectively. Structural figures were generated using PyMOL^18^ (version 2.5.2; Schrödinger).

### Phylogenetic and Residue Conservation Analysis

NCBI Protein BLAST^19,20^ (blastp) was used to search for homologous sequences using Nmar_1309, Nmar_0206, acetyl-CoA synthetase (ADP-forming) from *Candidatus* Korarchaeum cryptofilum, and Human Succinyl-CoA Synthetase (ADP-forming) as separate query templates (NCBI WP_012215692.1, WP_012214589.1, WP_012308855.1, and NP_003840.2, respectively). 45 sequences were selected based on species identifiers and used to construct a multiple sequence alignment (MSA) in MEGA (v11.0.13)^21^ with the default settings for CLUSTALW program (v2.1)^22^. This MSA was used to generate an amino acid-based phylogenetic tree of ACDs by the neighbor-joining method, using default settings within MEGA. The phylogenetic tree was then visualized with the iTOL: Interactive Tree of Life webserver^23^.

The Consurf web server ^24^ was used to estimate evolutionary conservation at each amino acid position with default parameters and automatic homolog selection. Interprosurf^25^ was used for interface area and hydrophobicity analysis.

## Results and Discussion

### Overall Homodimeric Nmar_1309 Structure

The 3-hydroxypropionyl-CoA synthetase Nmar_1309 is a homodimer consisting of CoA- and ATP-binding domains. These domains can be further subdivided into two CoA-binding subdomains (subdomains 1&2), two ATP-grasping domains (subdomains 3&4), and a CoA-ligase subdomain (subdomain 5). The sequences and overall structure are very similar to previously determined structures for other ACDs except for the characteristic domain shuffling resulting in a unique linker between the 4th and 1st domains and an overall subdomain sequence of [4-3-1-2-5](see Fig. 2). Subdomain 1 contains a Rossmann fold where the adenyl head of CoA binds in homologous structures. Subdomains 2 and 5 from opposite monomers interact to form a substrate binding pocket for 3HP and the sulfinyl-tail of CoA in addition to the reaction center itself. Two power helices contributed by subdomains 2 and 5 meet at their positive terminus at the reaction center and stabilize a phosphate bound there (Fig. 3) ^9^. Subdomain 2 also contains a catalytically important ‘swinging-loop’ (T493-S510) which bridges the 38Å between the ATP-grasping and the active sites and can be seen with a phosphate interacting with H503 at its tip (Fig. 3).

**Fig. 2:**
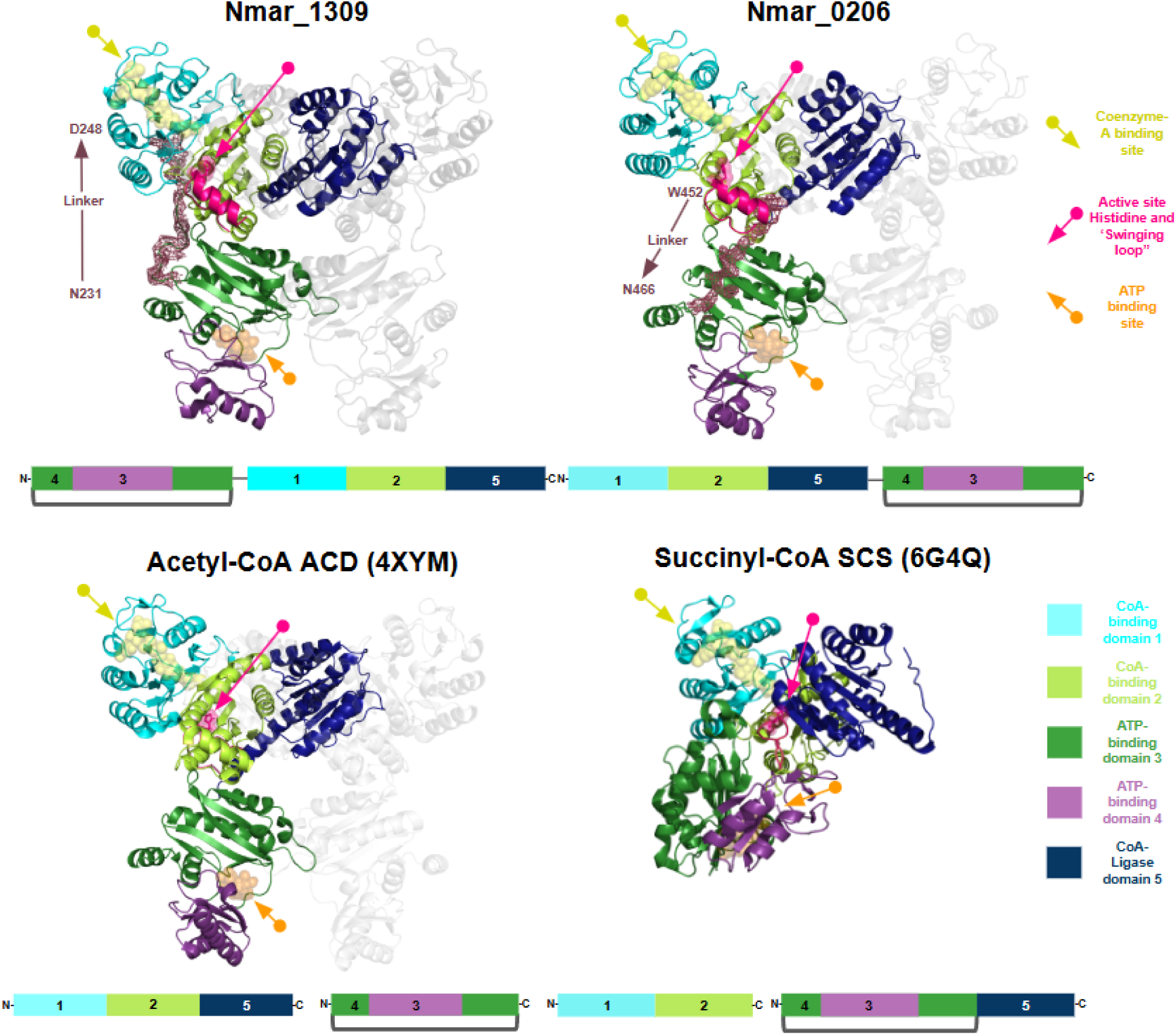
Domain organization comparison of Nmar_1309 protein with its homologs Nmar_0206, *Ca.* Korarchaeum cryptofilum acetyl-CoA synthetase and Human succinyl-CoA synthetase. Domain organization of domains 1 (teal), 2 (limon-green), 3-4 (forest-green and purple), and 5 (dark blue) are shown between homologs. Nmar_1309 and _0206 linker directionality is indicated by the arrow, along with starting and ending residues, and their respective densities are highlighted in red. Coenzyme-A (yellow) and ATP (orange) binding sites are also indicated by arrows, and their locations are shown by transparent blobs. The active site histidine and associated ‘swinging-loop’ is shown in pink, with the active site histidine pointed towards the active site.

**Fig. 3:**
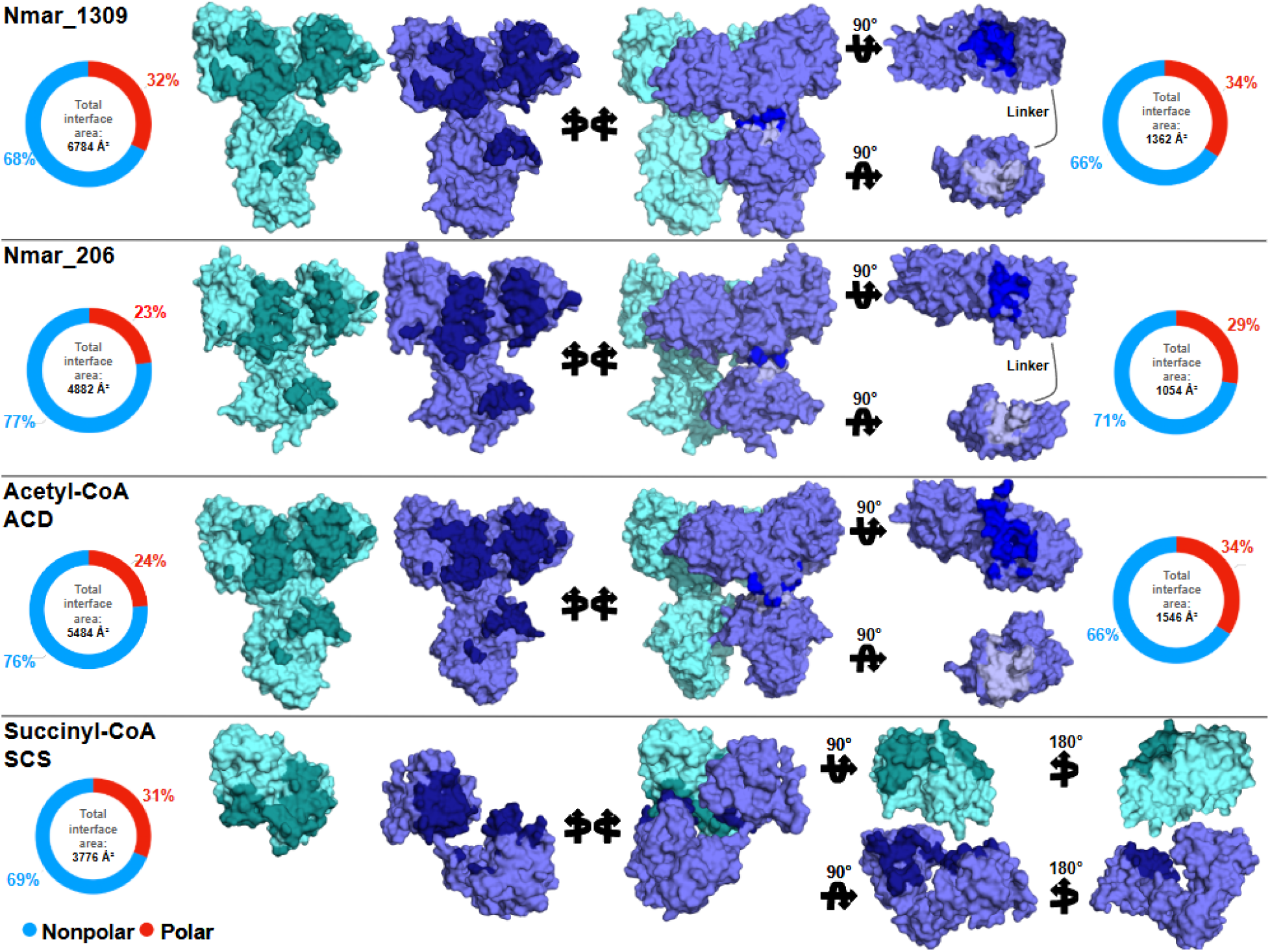
Comparison of interface surfaces of Nmar_1309, Nmar_0206, AC-S, and SCS. Chains A (teal) and B (slate) of Nmar_1309 and Nmar_0206 as well as the respective sites in AC-S and SCS are shown. Combined interface area between the Chains (left) as well as the ADP- and CoA-binding domains (right) are highlighted in differing shades. The pie charts show nonpolar (blue) and polar (red) surfaces.

Subdomains 3 and 4 are ATP-grasping domains with subdomain 3 being found within subdomain 4. The two chains of Nmar_1309 display conformational differences in subdomain 3 with chain A ‘closing’ 16° towards a bound ADPNP with residues P36 and K116 acting as hinge points. At the C-terminus of subdomain 3 is a 19 residue linker spanning from N231 to D248 connecting the ATP- and CoA-binding superdomains. This linkage is the basis for the homodimeric structure of Nmar_1309 and contrasts with the respective linker of Nmar_0206 and the lack of any linker within heterotetrameric and heterodimeric homologous structures (See Fig. 2).

### Domain Shuffling: Comparison with homologs Nmar_0206, acetyl-CoA synthetase and succinyl-CoA synthetase reveals structural differences

We have compared Nmar_1309 with its homologs Nmar_0206 (pdb:8WZU), Acetyl-CoA Synthetase (ADP-forming) from *Ca.* Korarchaeum cryptofilum (hereafter referred to as AC-S; pdb:4XYM) and Human Succinyl-CoA Synthetase (ADP-forming) (SCS; pdb:6G4Q) which share 36%, 38% and 22% sequence identity respectively. While Nmar_1309 has a domain organization [3-4-1-2-5], Nmar_0206 is organized [1-2-5-3-4], AC-S [alpha(1,2,5)/beta(3,4)] and SCS [alpha(1,2)/beta(3,4,5)] with alpha and beta referring to separate chains for the ATP- and CoA binding superdomains (see MSA in Supplementary Fig. 1). This domain shuffling between Nmar_1309 and its homologs is a common characteristic of the ACD superfamily. The observed domain shuffling suggests structural differences in the 3HP/4HB enzymes compared with their homologues and might give insights into their environmental variation in their evolutionary history. Previous studies have shown that other enzymes demonstrating shifts in domain organization present differing surface regions for regulation, change reaction chemistry, or even catalyze completely different reactions^10,26,27^.

The active site of Nmar_1309 lies at the interface between subdomains 1, 2, and 5 (Fig. 2). Subdomains 1 and 2 seem largely paired in homologous structures to maintain the active site and, as their N- and C-termini lie behind these subdomains (with respect to the active site), would therefore require a long linker to wrap around the structure to bring a directly bound subdomain 5 to that same active site for interaction (Supplementary Fig. 2). Citrate lyases seem to uniquely contain a large linker between these subdomains and display the monomeric form [x-1-2-3-4-5] (x being unique to these structures). To our knowledge no structure has been published to pdb with a resolved linker, so the exact orientation of these linkers are unknown. With citrate lyases as an exception, the structural limitations of directly linking subdomains 1, 2 and 5 in such a way that forms an active site seems evolutionarily constrained.

In Nmar_1309, a solvent-exposed linker region further contributes to structural integrity through interactions with subdomains 4 (N231, K232, K234, K235, N237, and S238), 2 (S240 and S245), and 1 (D248) (Supplementary Figure 3). These interactions may enhance the stability and dynamic communication between subdomains, potentially influencing its functional properties.

To further inspect the role of linkers between the CoA-binding and ATP-binding domains of Nmar_1309, we compared interdomain interfaces between homologues with different oligomeric forms. Nonpolar interactions are vital for the formation of functional dimeric enzymes^28^. While this was true for the four structures studied, variation in the hydrophobicity of the interface may speak to variability in the overall strength of the interfaces of these oligomers and, by extension, energetic environmental constraints. With regards to interfaces between the identical functional subunits (i.e., between chain A and B of Nmar_1309, and chains A/B and chains C/D of AC-S), relative levels of polar interactions between the four oligomeric forms were Nmar_1309 (32%) > SCS (31%) > AC-S (24%) > Nmar_0206 (23%) (see Fig. 3), with nonpolar interactions being the inverse. This may be explained in part by the relative surface area size of the Nmar vs AC-S structures, with the SCS structure being an outlier as the general curvature of the interface may support greater hydrophilicity.

The interfaces between CoA- and ATP-binding superdomains, however, tell a different story. Firstly, both Nmar structures have smaller interfaces than AC-S (1362 Å² and 1054 Å² vs 1546 Å²). Secondly, the AC-S structure has a much higher interface hydrophilicity between the CoA- and ATP-binding superdomains when compared to the interface between the chains (24% to 34%). While this trend is true for the Nmar interfaces (Nmar_1309: 32% to 34% and Nmar_0206: 23% to 29%), the relative increase in AC-S is much greater (10% increase vs 6% and 2%). As these interface areas are much smaller, this potentially suggests the need for the increased hydrophilicity with the linker itself reducing that need by stabilizing the structure without these bonds as previously proposed^3^.

### Phylogenetics suggest the divergence of 3HP from 4HB synthetases occurred following The Great Oxygenation Event

Phylogenetic analysis of homologous structures with different oligomerization states largely grouped into separate clades. There is a stark separation between citrate lyases and SCSs and other groups studied (Figure 4). This may be linked partly to the ability for citrate lyases and SCSs to form functional CoA lyase active sites from single functional units. Within the further delineated AC-S, 4HB, and 3HP synthetase clades, it has previously been noted that bacteria occupy an intermediate position between AC-Ss and the other clades, suggesting horizontal gene transfer^3^. This is further supported by the findings that a high number of genes within thaumarchaeal genomes appear to originate in Bacteria (instead of Archaea)^29^.

**Fig. 4:**
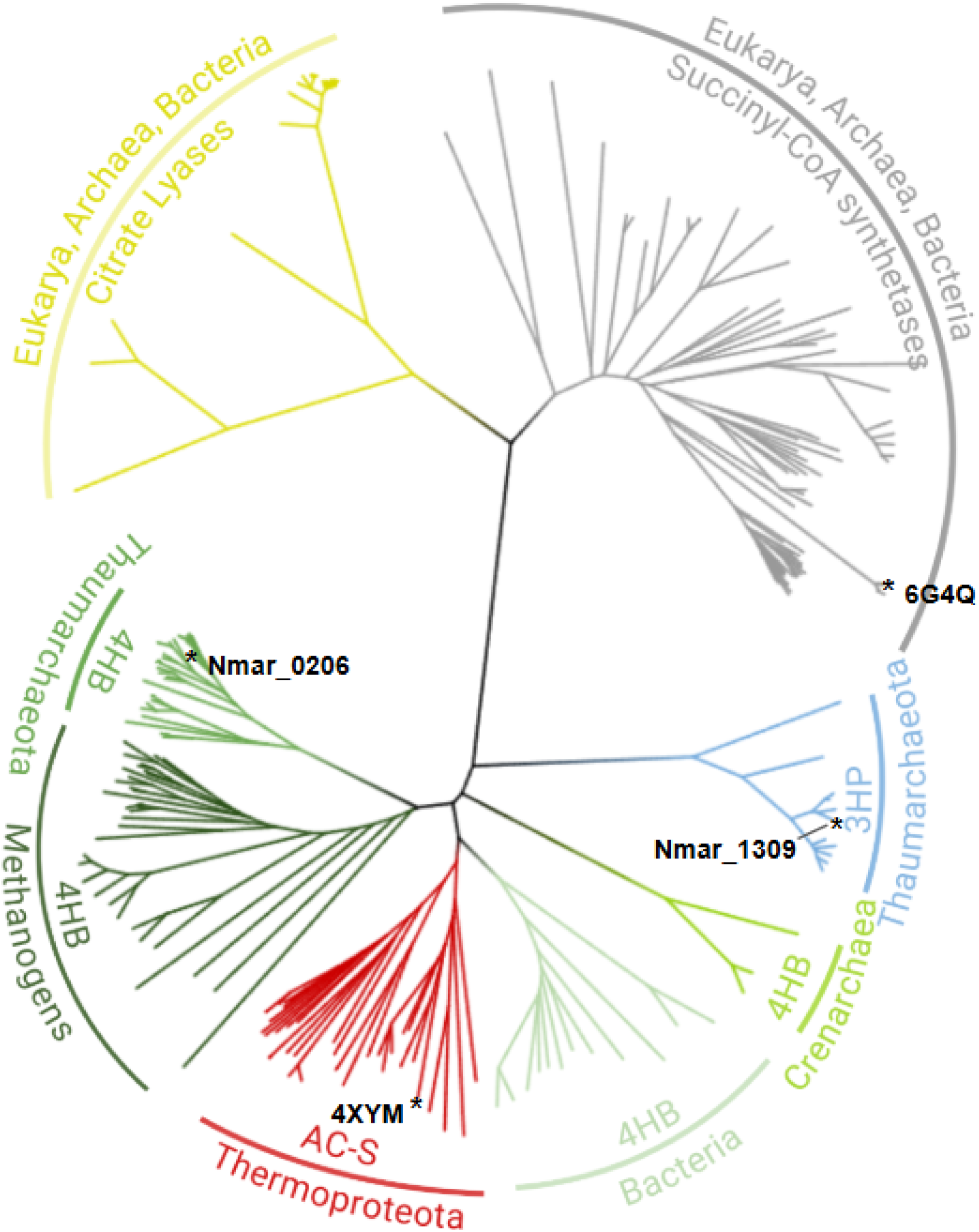
Phylogenetic tree of Nmar_1309 and homologues. This tree highlights the strong divergence between enzymes with different oligomerization structures. Heterodimeric SCSs and homohexameric Citrate Lyase structures with the domain organization x-1-2-3-4-5 (x being unique to these enzymes) makeup a clade separate from heterotetrameric AC-Ss and homodimeric 3HP and 4HB synthetases. While citrate lyases and SCSs are present in Eukarya (not separated for simplicity), none of the other groups are found in Eukarya. Asterisks (*) indicate structures that were compared in Figure 2 (i.e., Thaumarchaeota Nmar_0206 4HB and Nmar_1309 3HP, Thermoproteota Acetyl-CoA 4XYM AC-S, and Succinyl-CoA Synthetase SCS 6G4Q).

A previously described evolutionary timeline mapped thaumarchaeal divergence from crenarchaeota and euryarchaeota to the great oxygenation event^29^. Interestingly, a few crenarchaeal 4HB synthetase homologues analyzed here formed their own clade between Thaumarchaeotal 3HP and 4HB synthetases (Figure 4). This includes the model organism *M. sedula* used to originally describe the 3HP/4HB cycle^6^ but which generally uses the less energetically efficient AMP-forming versions of these enzymes^2,6^. The fact that crenarchaeal homologues clustered between the two thaumarchaeal enzymes suggest that the divergence between the 3HP and 4HB enzymes in Thaumarchaeota may have happened following the oxygenation of the oceans^29^. The close homology of Thermoproteota AC-S and thaumarchaeal 4HB synthetases may indicate that these evolved earlier in the evolutionary timeline.

### Structural Comparison of Chain A and Chain B of Nmar_1309 demonstrates conformation differences in the ATP-binding domain

The Nmar_1309 structure contains 3 ligands and 2 bound phosphate molecules functionally relevant to the reaction mechanism. Within each active site at the juncture of the CoA-binding and ligase-CoA domains, 3HP can be found stabilized by the presence of a phosphate at the head of two ‘powerhelices’ (see Fig. 5a,b)^9^. Furthermore, a single ADPNP is bound in the ATP-binding domain of chain A but not chain B (see Fig. 5c-d). The well defined electron densities in this region display conformational differences essential for the binding and stabilization of ATP before dephosphorylation (see Fig. 5e-f; Fig. 6, supplementary Fig. 4). When the monomers are aligned, a ∼15.6° rotation of the ATP-grasping domain of chain A can be seen closing towards the bound ADPNP (averaged across residues H69, R96 and K88 with M111 as a stable reference; see Fig. 5g and Supplementary Fig. 5). In contrast, Chain B, shows an electron density map without the ligand displaying a more open conformation rotated away from the binding site.

**Fig. 5:**
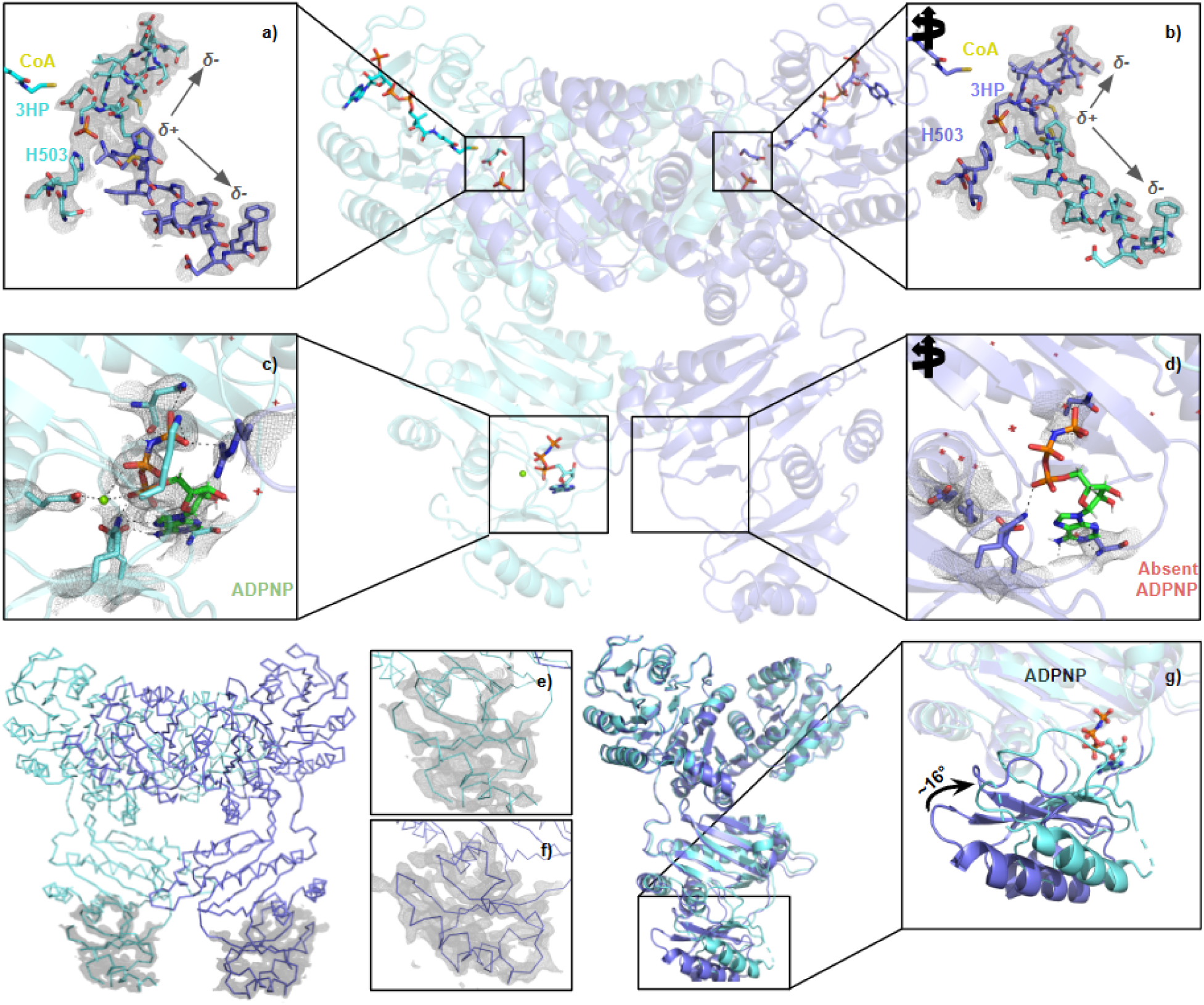
Bound ligand densities and ATP-binding domain conformational shifts. Structural localization of 3HP and ADPNP binding are shown here, along with their respective electron densities. **a,b** The active site is found at the head of two power helices, which stabilize the negative charge of the bound phosphate. **c** In Chain A, the ADPNP binding site and its electron density are shown, **d** while in Chain B, the electron density shows a lack of ADPNP (the one shown is placed at the relative location from a superposed chain A) at that same site. **e,f** The electron densities of the two chains’ ATP-binding domains. **g** A structural alignment of Chain A and Chain B illustrates a ∼16° rotation upon ADPNP bonding. The rotation was calculated by averaging the angles formed between the locally stable M111 residue and the shifted residues H69, R96, and K88. 2Fo-Fc electron density maps are shown with a contouring level of 1σ and carved to 2 Å.

**Fig. 6:**
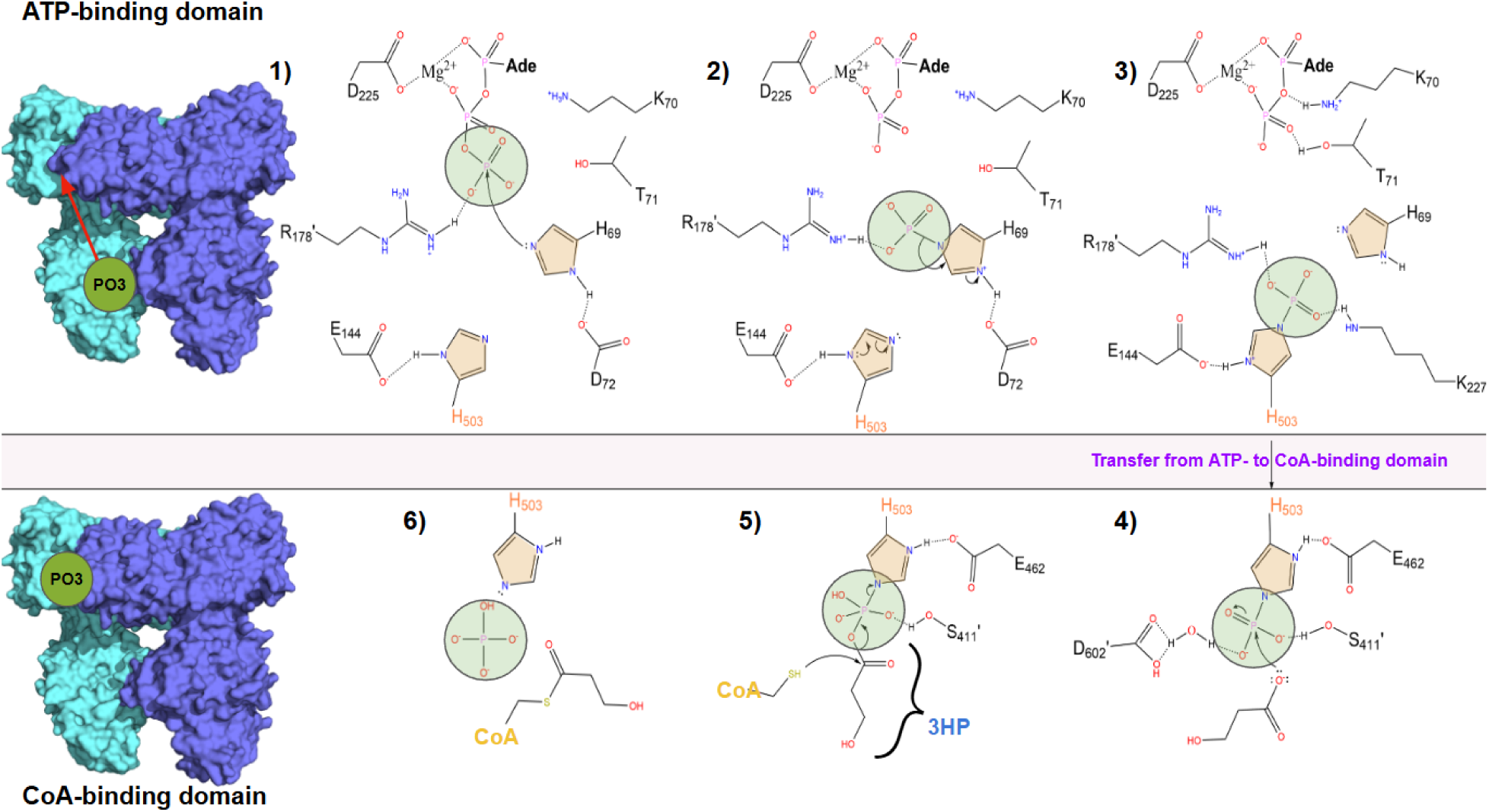
Reaction mechanism of Nmar_1309. **1-6** The proposed reaction mechanism of Nmar_1309 derived from homologous structures. Following ATP binding, the gamma phosphate of the ATP is dephosphorylate and passed from **1,2** H69 to **2,3** H503 on the ‘swinging-loop’ of subdomain 2 in the CoA-binding domain. **4,5** The ‘swinging-loop’ subsequently swings into the CoA-binding domain active site and is stabilized by Asp602’ and Ser411’, where it may interact with the acyl-group on 3-HP. This leads to the formation of a transient 3HP-phosphohistidine before reaction with CoA. **6** shows the products of the reaction including the free phosphate, unbound H503 and 3-hydroxypropionyl-CoA. Surface representation of the structure on the left highlights the 41Å phosphoryl transfer between domains.

### Observed binding modes of 3HP and phosphates in Nmar_1309 offer insight into reaction mechanisms

The crystal structure reveals two 3HP molecules, one bound at each active site to H503 and a phosphate molecule. Due to the low resolution of the model, two opposite-facing conformations of each 3HP were placed within the density and occupancy refined (See Supplementary Fig. 6). Our results show that the 3HP molecules favor the binding orientation seen in homologous proteins (i.e., carboxylate end oriented towards the phosphate refines to 70% occupancy in chain A and 74% in chain B; see Supplementary Fig. 7)^12,30,31^ and are suggested by the reaction mechanism itself (See Fig. 6).

### Reaction mechanism of Nmar_1309 and structural localization

The reaction mechanism for ACDs has been established by several previous studies^9,10,11,12,30,31,32,33,34,35^. The larger distance between the ATP-binding and the CoA-binding active sites and the presence of an additional histidine in the ATP-binding site support the 2 histidine reaction mechanism^10^. This is one of two mechanisms known for ACDs which have been described distinguishing SCSs, which have an ATP binding domain closer to the CoA binding site (and therefore the reaction site), from those further apart like Nmar_1309. SCS originally described a reaction mechanism involving a single histidine on the aforementioned swinging loop^9^ whereas subsequently ACDs with larger distances between the two binding sites have an additional histidine in the ATP binding site^10^. All residues involved have been inferred by sequence alignment and from their respective locations upon superposition.

The reaction begins with the binding and dephosphorylation of the gamma phosphate of ATP by H69 within the ATP binding pocket^9,11,32,33^. The removed phosphate is initially covalently bound to H69 but is subsequently passed to H503 which, attached to the flexible ‘swinging loop’, swings into the reaction center between subdomains 1, 2 and 5 upon phosphorylation^11,12,31,33^. Stabilized by the positively charged dipole of the power helices and potential interactions with other neighboring residues (such as E462 and D602’), at this point, phosphoHis503 is available for interaction with 3HP^12,31^. Incoming 3HP covalently binds to phosphoHis503 forming a transient 3HP-phosphoHis503 ^35^ leaving it susceptible to nucleophilic attack by CoA to peel off the 3HP, freeing phosphate and forming 3HP-CoA^12^.

Although unphosphorylated, this structure of Nmar_1309 appears to be temporally located similar to a phosphorylated H503 in the CoA-binding active site interacts with 3HP just before the 3HP-phosphoHis503 intermediate formation (see Fig. 6, step 4). Both active sites show interactions predicted at that step between D602’, Phosphate, H503 and a bound water (with the exception of a single bond in chain A and the obvious lack of a covalent bond between phosphate and histidine; see Fig. 7). This being said, many interactions which can be seen were not reported in homologous structures including with N376, G558’, A557’, G412, T413, N555, D636’, V434 and additional waters. Most of these interactions are shared between the two active sites with exceptions in the single additional D602’; the additional interactions with 3HP, V434 and an intermediate water; and those of the additional water between 3HP, phosphate, N555 and D602’. While these interactions may just be a product of the bound phosphate instead of phosphohistidine at this reactive step, they may reflect similar interactions in the real reaction mechanism thus far unelucidated.

**Fig. 7:**
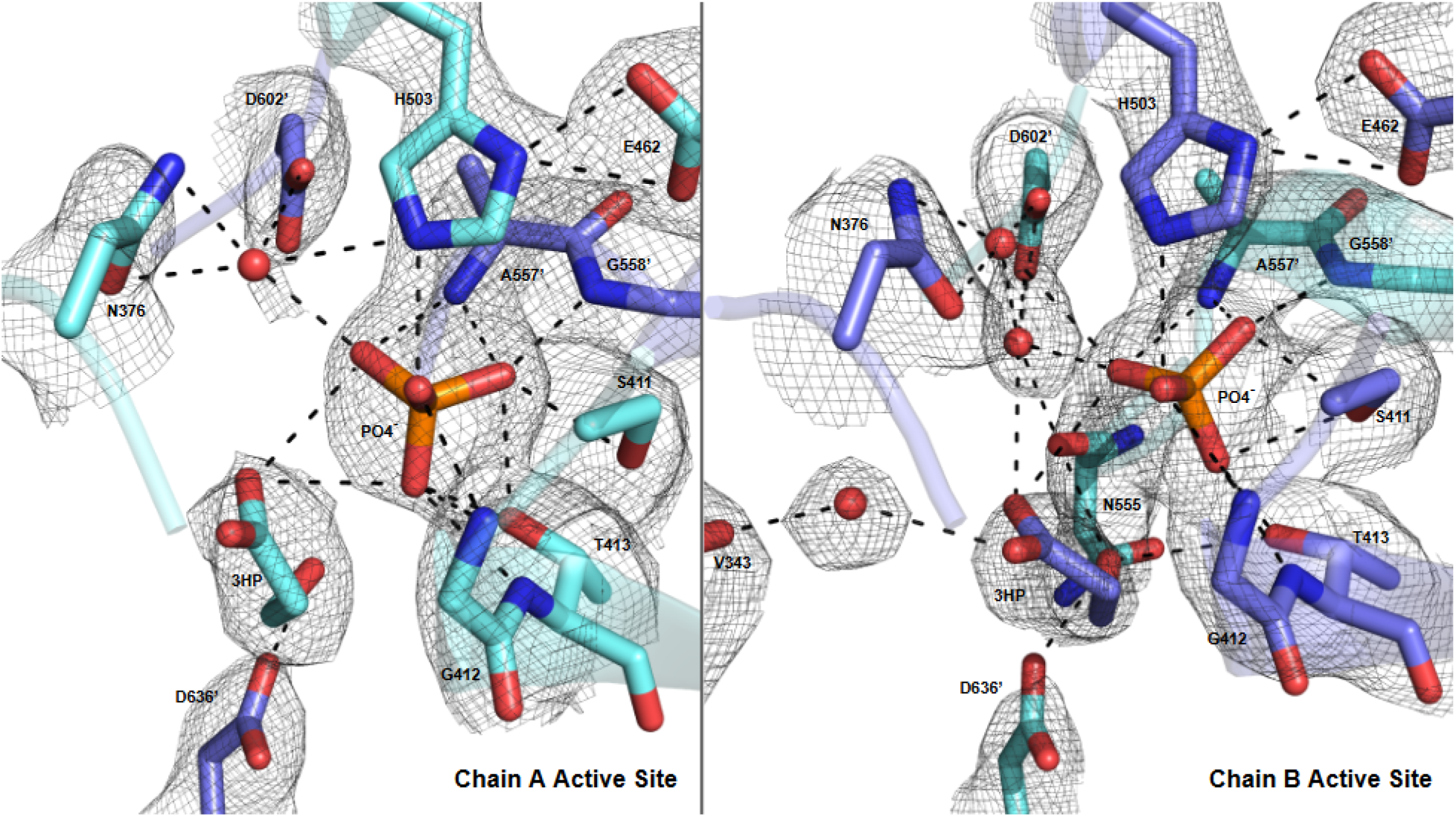
Active Site interactions with bound phosphate and 3-hydroxypropionate. A free phosphate can be found at the active site bound to H503 and the ligand of interest 3-hydroxypropionate. Most interactions here are not found in the predicted reaction mechanism with the exception of those between E462 and H503; between S411 and phosphate; and between D602’, a water molecule and phosphate seen in Figure 6 panel 4.

## Conclusion

The efficiency of global carbon fixation is driven by diverse enzymatic pathways, with the thaumarchaeal 3HP/4HB cycle standing out for its energetic optimization. Our analysis of Nmar_1309 reveals key evolutionary and structural adaptations that underpin this cycle’s success. The evolution of Nmar_1309 seems tied to the evolution of thaumarchaeota, with its divergence from Nmar_0206 and other ACDs potentially a later development in evolutionary history. Domain shuffling is characteristic of ACDs and allows evolutionary flexibility in how these structures are organized resulting in a number of oligomeric types (see Fig. 2). Linkers between chains play an important role in stabilizing oligomerization and active site formation (see Fig. 3). A large physical distance between CoA-binding and CoA-lyase subdomains in a functional active site makes a direct linker between these a potential evolutionary barrier (see Fig. 4; see Supplementary Fig. 2). AC-Ss, 4HBSs, and 3HPSs seem to overcome this by oligomerizing. Citrate lyases, in contrast, do form this linker while SCSs form a heterodimer instead. Due to the evolutionary distance between these oligomerization states, this may suggest the evolutionary time required to transition to functional active sites in single asymmetric units. Nmar_1309 and Nmar_0206 both have linkers between CoA-binding and ATP-binding domains but link on opposite ends of subdomain 3. It is worth noting that while many other archaea - particularly representatives from thermoproteota and crenarchaeota - contain both heterotetrameric and homodimeric forms of ACDs (4HB synthetases and AC-Ss), *Nitrospumilus* sp. do not. This could suggest the evolutionary benefit of a linker connecting CoA-binding and ATP-binding domains in oxygenic environments that formed during the evolution of thaumarchaeota.

Many standard features of ACDs can be seen in the structure of Nmar_1309 including the catalytically important power helices, locally stabilized phosphate and bound 3-hydroxypropionate (3HP) at both active sites (see Fig. 5). Both active sites indicate very similar binding modes, suggesting a temporal stabilization near step 4 of the reaction mechanism while displaying many interactions not previously discussed (see Fig. 6 and 7). Whether these are a product of the free phosphate or reflect catalytically important interactions need further investigation.

This structural elucidation of Nmar_1309, particularly when combined with the structure of Nmar_0206, completes the characterization of key ADP-forming Acyl-CoA synthetases in the vital thaumarchaeal 3HP/4HB cycle. Considering the significant contribution of Thaumarchaeota to global carbon fixation, unraveling the structural and evolutionary intricacies of their metabolic enzymes—products of extensive adaptation and inherently sensitive to environmental shifts—is of profound importance. The detailed insights gained from such studies are fundamental not only for understanding natural biogeochemical processes but also for providing a robust blueprint for innovative bioengineering strategies to address pressing global challenges.

## Data Availability Statement

The Nmar_1309 structure and associated files were deposited in the PDB under 9Y43. All other supporting data are available from the corresponding author upon reasonable request.

## Permission to reproduce material from other sources

All content, figures, and tables presented in this manuscript are the original work of the authors. No material has been reproduced from other sources.

## Contributions

H.D., Y.Y., C.A.F, S.W. conceived the study; H.D., J.J. designed the experiments; H.D., J.J., B.B.T., B.T., S.Y., M.Y., T.D. performed the experiments with input from S.W.. J.J. and B.B.T. performed phylogenetic analyses. All the authors took part in the interpretation of results and preparation of the manuscript.

## Acknowledgements

H.D. acknowledges support from NSF Science and Technology Center grant NSF-1231306 (Biology with X-ray Lasers, BioXFEL). This publication has been produced benefiting from the 2232 International Fellowship for Outstanding Researchers Program and the 1001 Scientific and Technological Research Projects Funding Program of the TÜBİTAK (Project Nos. 118C270 and 120Z520). However, the entire responsibility of the publication belongs to the authors of the publication. The financial support received from TÜBİTAK does not mean that the content of the publication is approved in a scientific sense by TÜBİTAK. The work conducted by the U.S. Department of Energy Joint Genome Institute (https://ror.org/04xm1d337), a DOE Office of Science User Facility, is supported by the Office of Science of the U.S. Department of Energy operated under Contract No. DE-AC02-05CH11231. SW and CAF acknowledge support from the U.S. Department of Energy (DOE) Office of Science, Biological and Environmental Research; Stanford Precourt Institute; and SLAC Laboratory Directed Research and Development. SSRL is supported by the U.S. Department of Energy (DOE), Office of Science, Office of Basic Energy Sciences (OBES) under Contract No. DE-AC02-76SF00515. The SSRL Structural Molecular Biology Program is supported by the DOE Office of Biological and Environmental Research and by the National Institutes of Health, National Institute of General Medical Sciences (NIGMS) (including P41GM103393).https://docs.google.com/document/d/15DKqmvS4hWirc7UkWO6nGVCpnu6zwXQnF-opz30YEp0/edit.

## Supplementary Figures

**Supplementary Figure 1.**
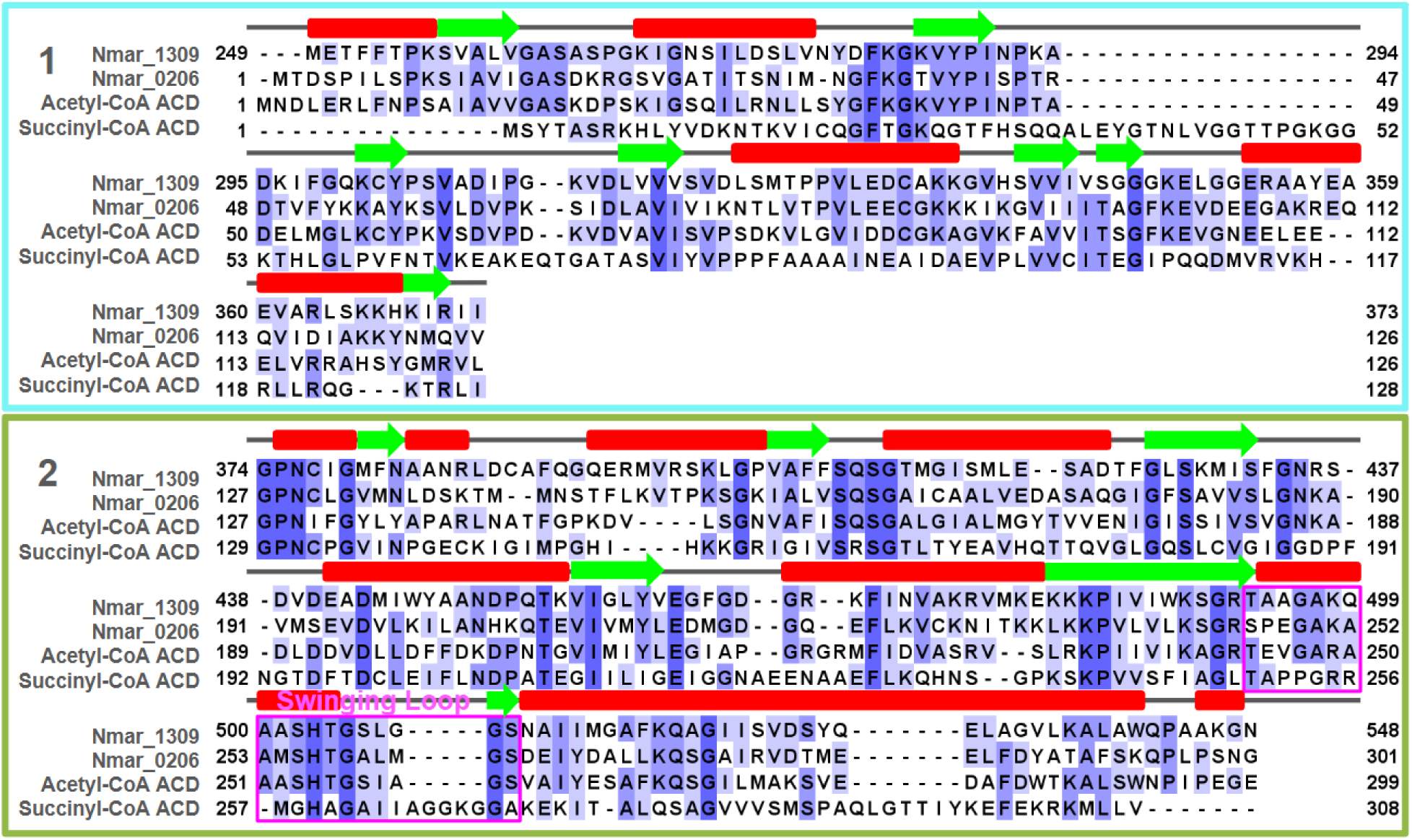
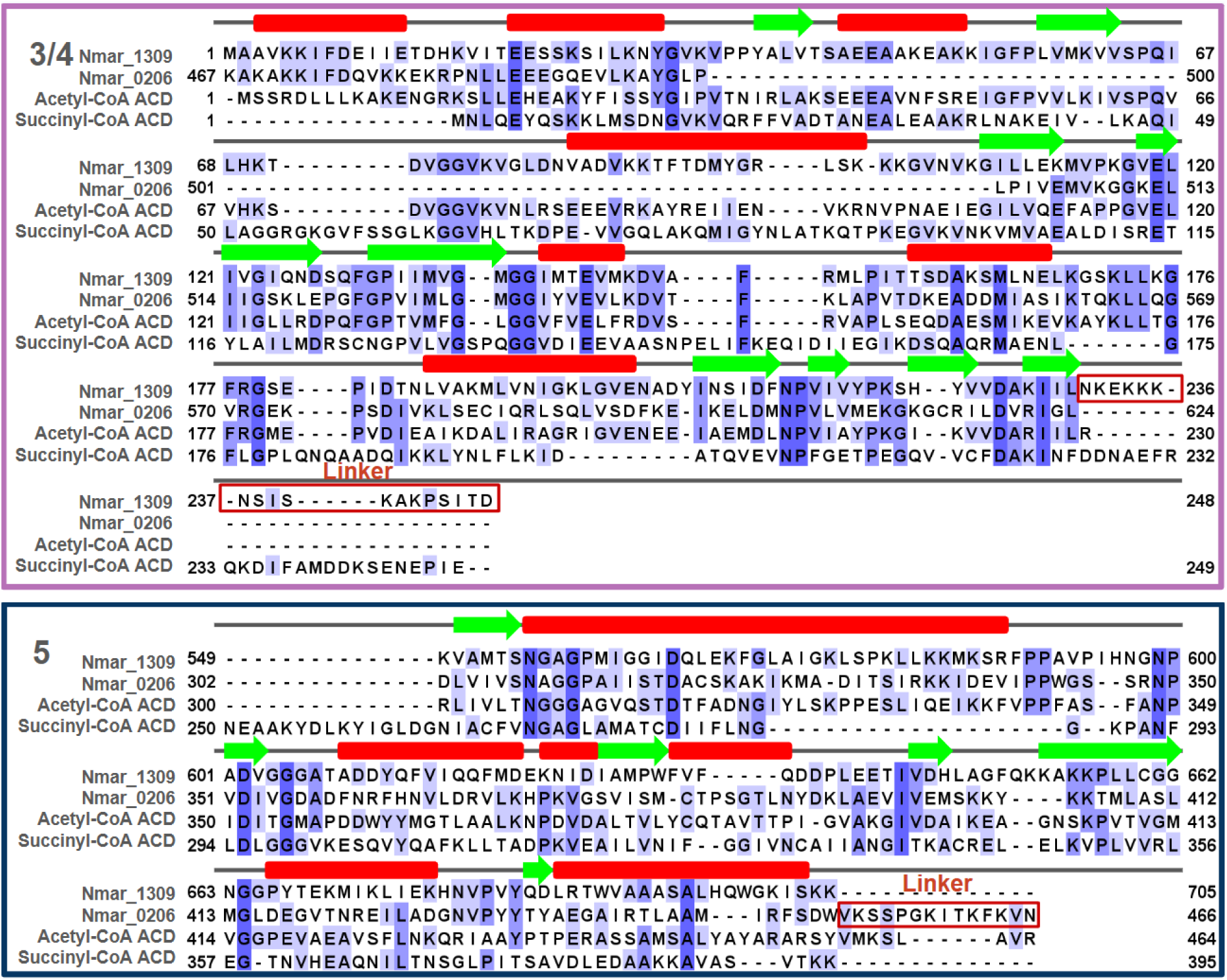
Domain sequence alignment of ACD homologues. Multiple sequence alignments for subdomains 1-5, showing residue and secondary structure conservation with beta-sheets indicated by green arrows and alpha-helices indicated with red bars.

**Supplementary Figure 2.**
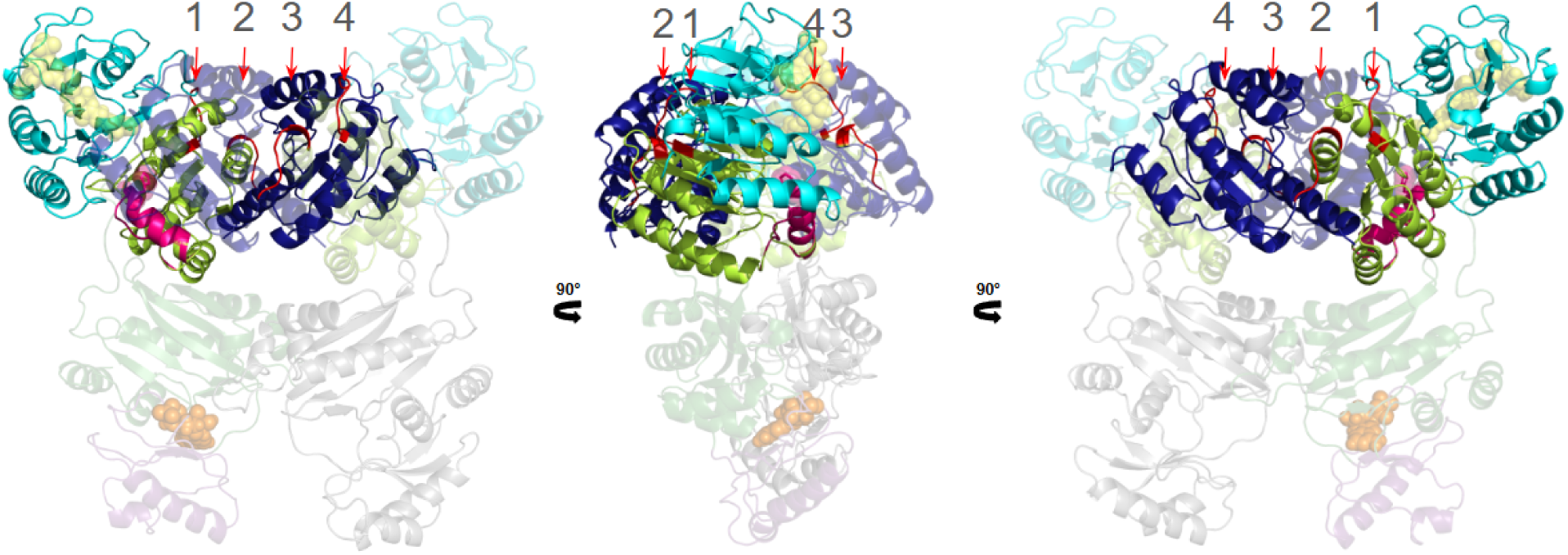
Electron density, residues and interactions of linker loop. Numbers 1 and 2 indicate connections between subdomains 1-2 and 2-5 on Chain A. Chain B, shown transparent in the figure, contains these same connections labeled numbers 4 and 3.

**Supplementary Figure 3.**
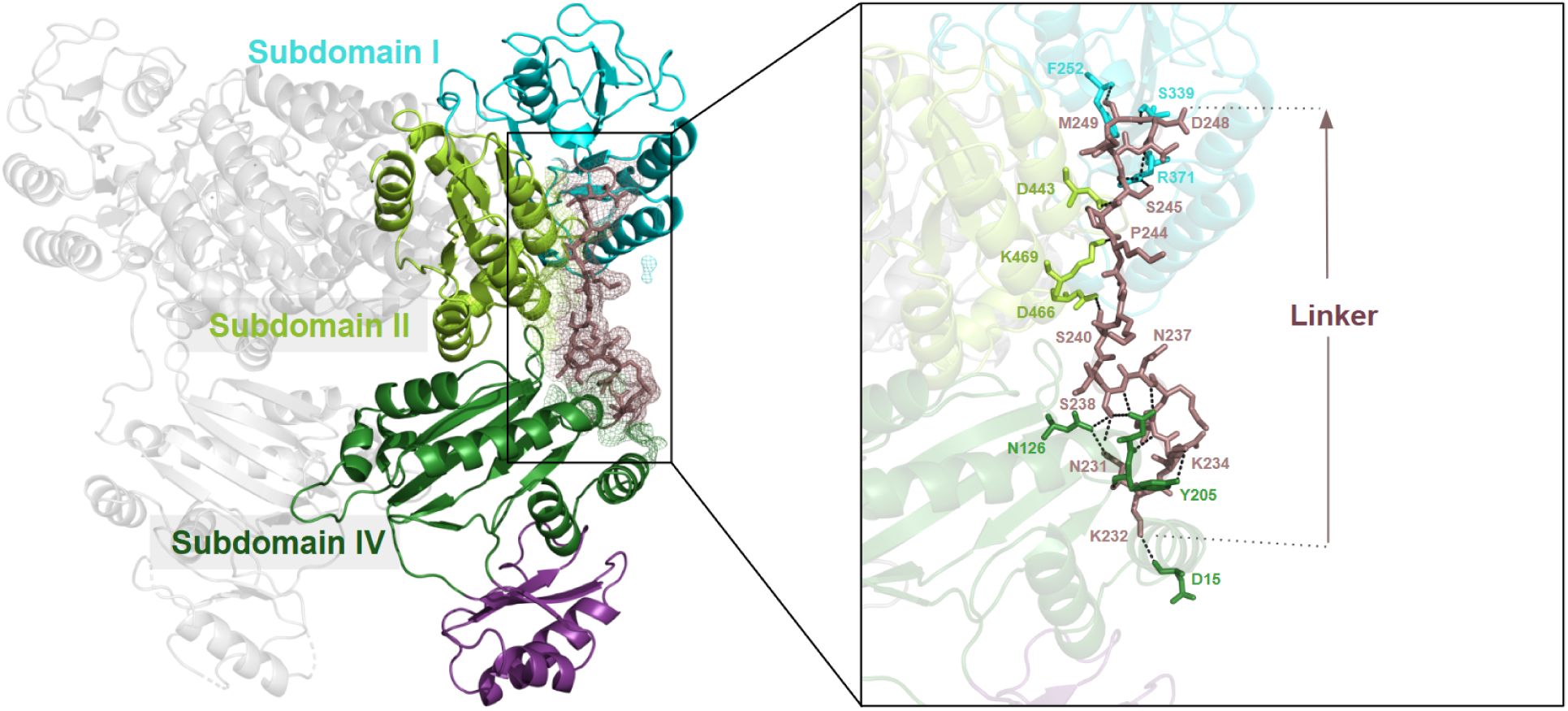
Interactions of the linker region with subdomains I, II, and IV. Subdomain I (teal), Subdomain II (lemon), and Subdomain IV (forest), along with the linker region (dirty violet) are shown in Chain A. The residues involved in interactions between the subdomains and the linker region are highlighted.

**Supplementary Figure 4.**
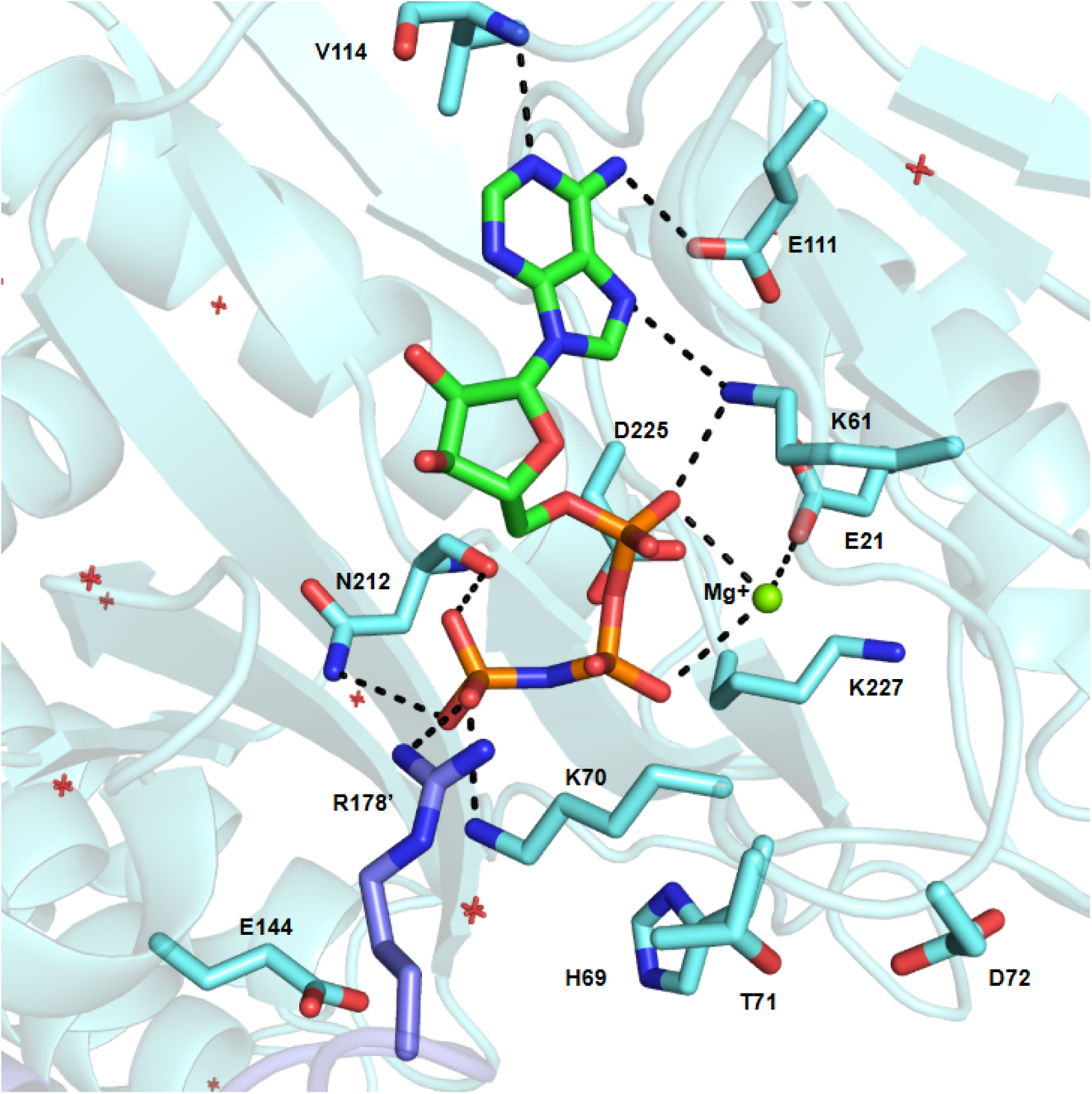
Bound ADPNP and Magnesium and their interactions within the ATP binding domain. ADPNP with a bound Magnesium ion can be found bound to the ATP binding site of chain A. While the interaction between the gamma phosphate of ADPNP and R178’ is consistent with the expected reaction mechanism (see Fig. 6 panel 1), other interactions are either missing (in the case of interactions between ADPNP and D225 and T71, H69 and D72) or interact differently than expected (K70 and its interaction with the gamma phosphate of ADPNP instead of the bridging oxygen between the alpha and beta phosphates). Additionally, interactions can be seen below the adenyl head include that of K61, E21 and N212. Interactions with the adenyl head (V114, E111 and K61) likely do not play a role in catalysis of the reaction itself, instead stabilizing the structure of ADPNP itself.

**Supplementary Figure 5.**
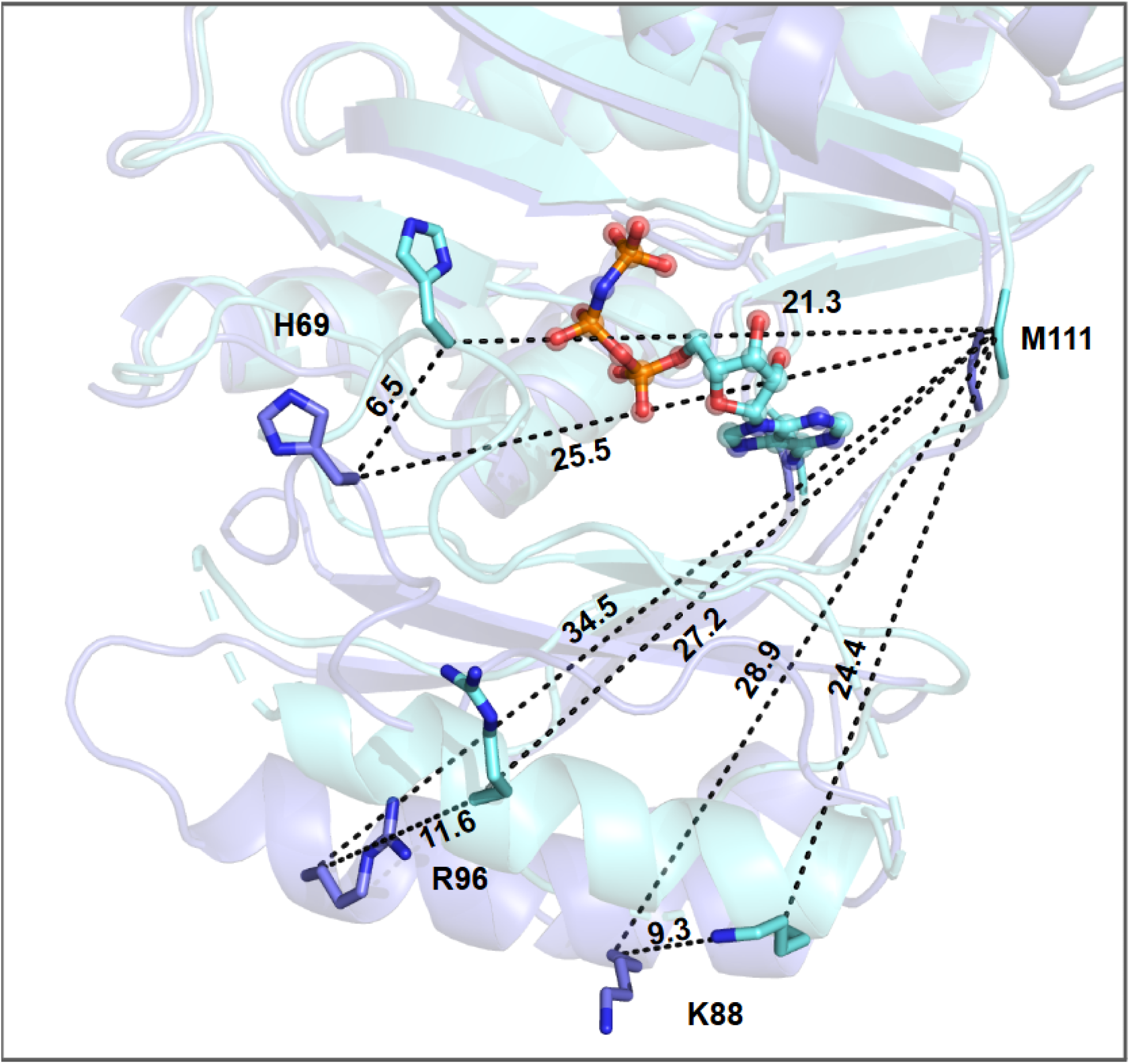
Calculating the rotation of ADP-grasping domain. The distances used to calculate ATP-grasping rotation upon ADPNP binding are shown. With M111 used as a reference, the average translation of residues H69, K88 and R96 were used to calculate that angle with respect to M111. These were averaged to achieve a rotation of 15.6°.

**Supplementary Figure 6.**
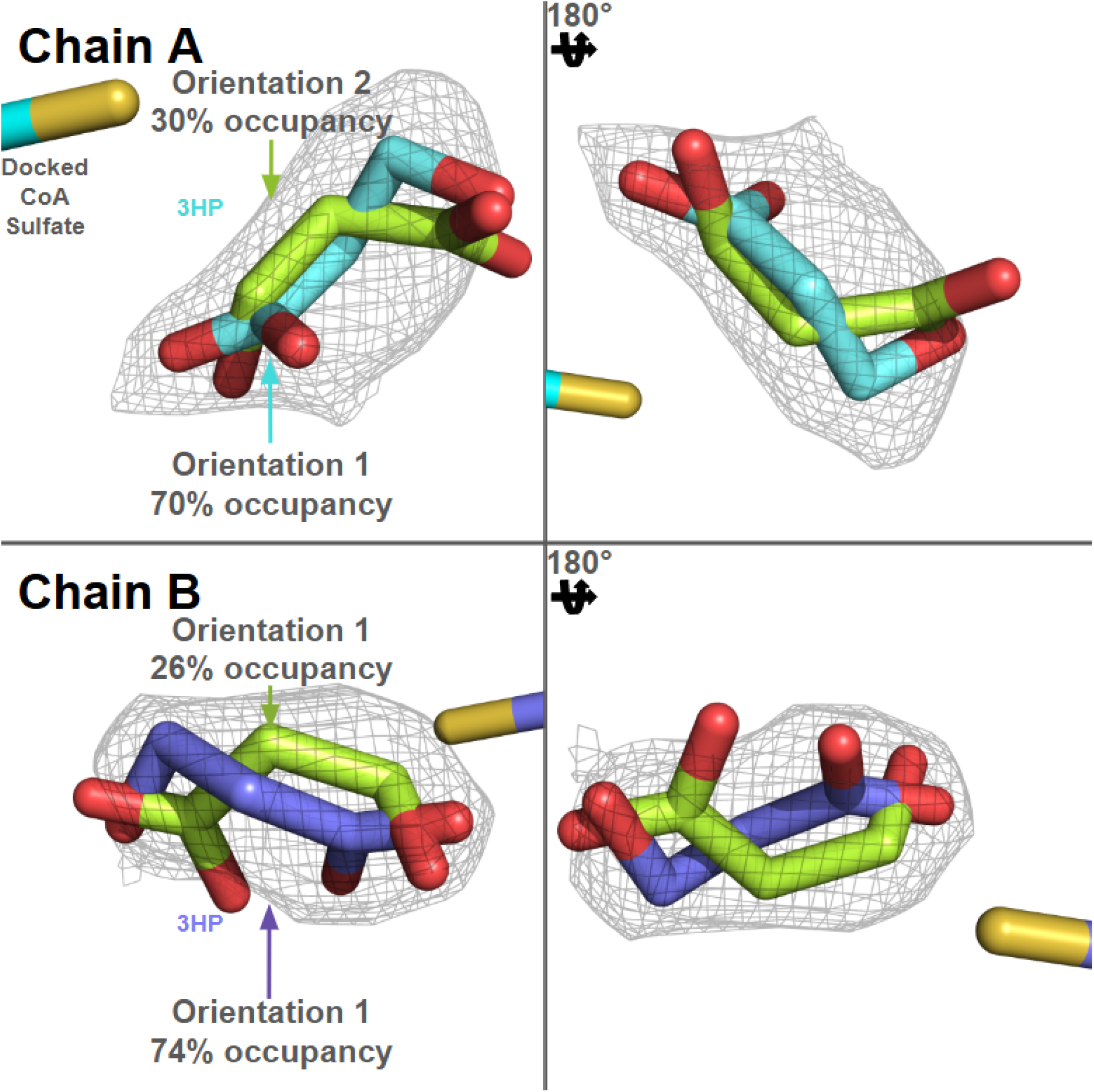
Occupancies of oppositely orentiented 3-hydroxypropionate in a low resolution electron density. Electron density and occupancy of the two possible orientations of 3HP from chain A and B. The dominant orientation seems to mirror other bound ligands in homologues. This orientation was used for the final figure generation. The sulfinyl tail of Coenzyme-A from PDB-ID 4YAK is shown for orientation.

**Supplementary Figure 7.**
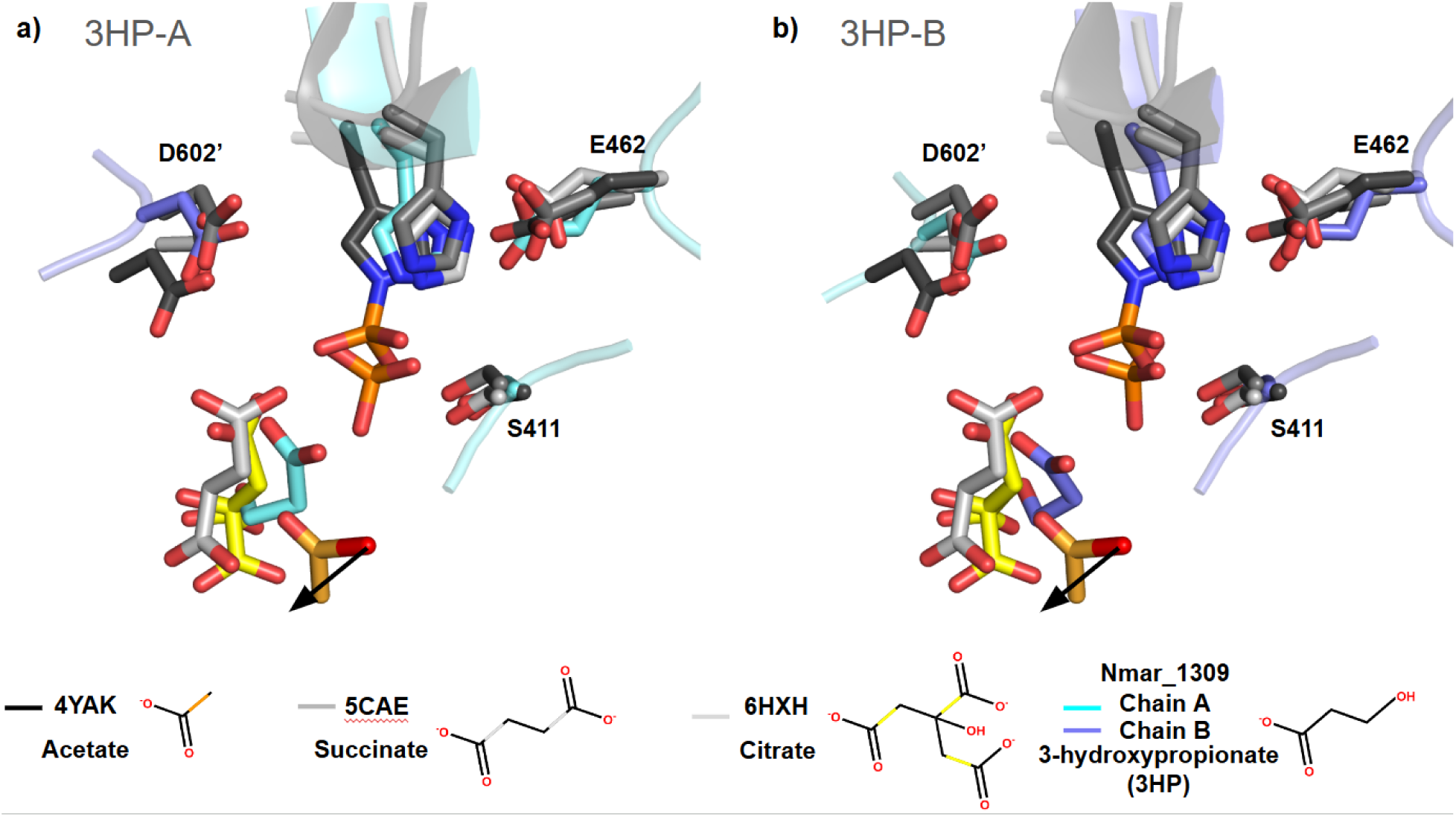
Active site comparisons of ACD homologous structures. Dominant 3HP occupancy orientations for 3HP within chains A and B can be seen here relative to AC-S, SCS and ATP Citrate Lyase bound ligands and their surrounding residues. The black arrows indicate the direction Coenzyme-A passes from the active site. The shown phosphates are those bound to this Nmar_1309 structure. All homologous ligands are bound to their respective structure with the exception of the acetate, which is found bound to the sulfinyl tail of Coenzyme-A in the pdb structure 4YAK.

**Supplementary Figure 8.**
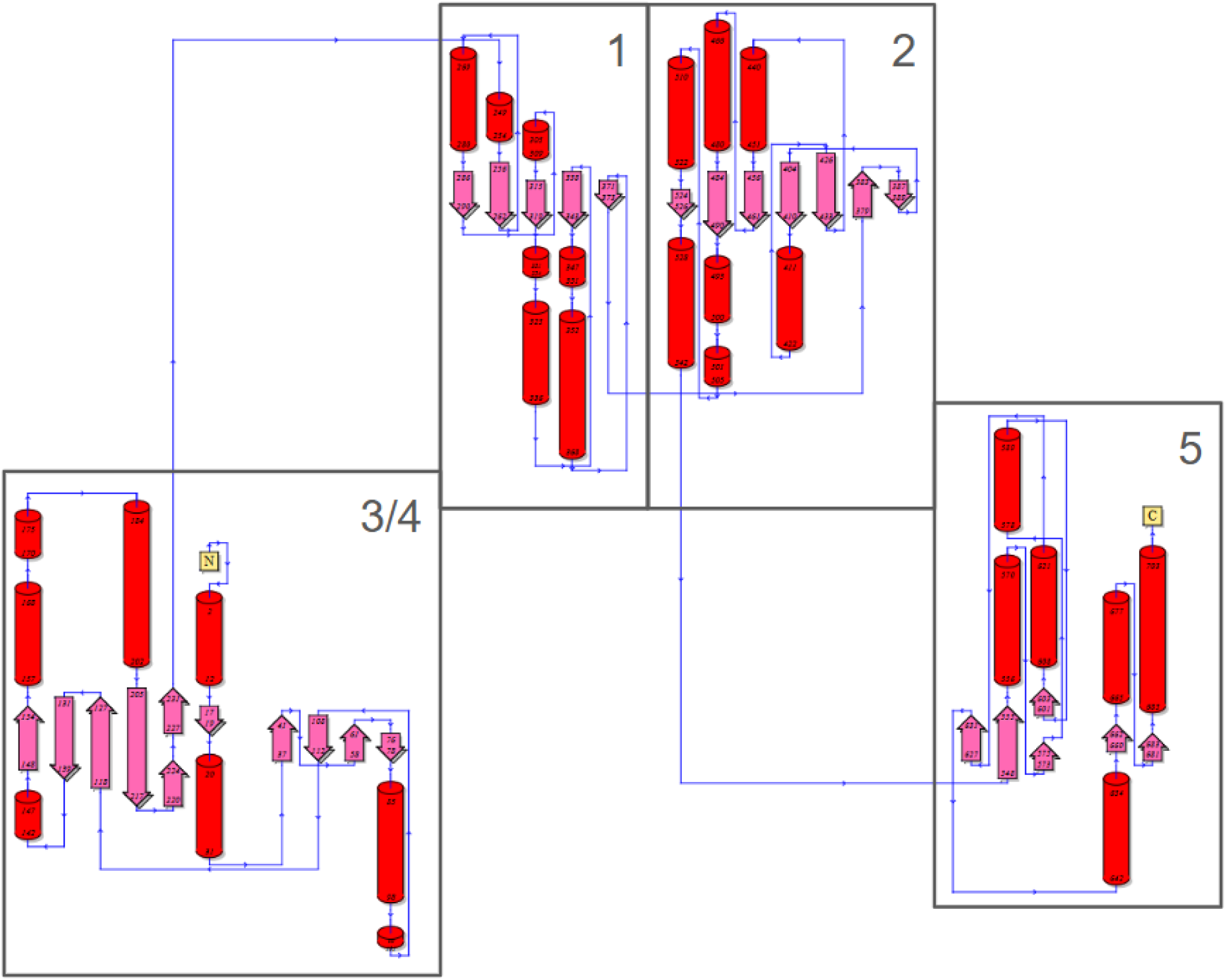
TOPS diagram of Nmar_1309. The TOPS diagram of Nmar_1309 shows the internal organization of the individual domains.

## References

1. World Meteorological Organization. (2024). State of the Global Climate 2023. https://library.wmo.int/records/item/68835-state-of-the-global-climate-2023

2. Könneke, M., Schubert, D. M., Brown, P. C., Hügler, M., Standfest, S., Schwander, T., Schada von Borzyskowski, L., Erb, T. J., Stahl, D. A., & Berg, I. A. (2014). Ammonia-oxidizing archaea use the most energy-efficient aerobic pathway for CO2 fixation. Proceedings of the National Academy of Sciences, 111(22), 8239–8244. 10.1073/pnas.1402028111

3. Johnson, J., Tolar, B. B., Tosun, B., Yoshikuni, Y., Francis, C. A., Wakatsuki, S., & DeMirci, H. (2024). Crystal structure of the 4-hydroxybutyryl-CoA synthetase (ADP-forming) from *Nitrosopumilus maritimus*. Communications Biology, 7(1), Article 32. 10.1038/s42003-023-05705-z

4. Destan, E., Johnson, J., Yoshikuni, Y., & DeMirci, H. (2021). Structural insights into bifunctional thaumarchaeal crotonyl-CoA hydratase and 3-hydroxypropionyl-CoA dehydratase from *Nitrosopumilus maritimus*. Scientific Reports, 11, Article 22941. 10.1038/s41598-021-02180-8

5. DeMirci, H., Johnson, J., Destan, E., Tolar, B. B., Yoshikuni, Y., Francis, C. A., & Wakatsuki, S. (2020). Structural adaptation of oxygen tolerance in 4-hydroxybutyrl-CoA dehydratase, a key enzyme of archaeal carbon fixation [Preprint]. bioRxiv. 10.1101/2020.02.05.935528

6. Berg, I. A., Kockelkorn, D., Buckel, W., & Fuchs, G. (2007). A 3-hydroxypropionate/4-hydroxybutyrate autotrophic carbon dioxide assimilation pathway in archaea. Science, 318(5857), 1782–1786. 10.1126/science.1149466

7. Ingalls, A. E., Shah, S. R., Hansman, R. L., Aluwihare, L. I., Santos, G. M., Druffel, E. R., & Pearson, A. (2006). Quantifying archaeal community autotrophy in the mesopelagic ocean using natural radiocarbon. Proceedings of the National Academy of Sciences, 103(17), 6442–6447. 10.1073/pnas.0510157103

8. Sánchez, L. B., Galperin, M. Y., & Müller, M. (2000). Acetyl-CoA synthetase from the amitochondriate eukaryote *Giardia lamblia* belongs to the newly recognized superfamily of acyl-CoA synthetases (nucleoside diphosphate-forming). Journal of Biological Chemistry, 275(8), 5794–5803. 10.1074/jbc.275.8.5794

9. Wolodko, W. T., Fraser, M. E., James, M. N., & Bridger, W. A. (1994). The crystal structure of succinyl-CoA synthetase from *Escherichia coli* at 2.5-A resolution. Journal of Biological Chemistry, 269(14), 10883–10890. 10.1016/s0021-9258(17)34141-8

10. Bräsen, C., Schmidt, M., Grötzinger, J., & Schönheit, P. (2008). Reaction mechanism and structural model of ADP-forming Acetyl-CoA synthetase from the hyperthermophilic archaeon *Pyrococcus furiosus*: Evidence for a second active site histidine residue. Journal of Biological Chemistry, 283(22), 15409–15418. 10.1074/jbc.m708767200

11. Fraser, M. E., James, M. N. G., Bridger, W. A., & Wolodko, W. T. (1999). A detailed structural description of *Escherichia coli* succinyl-CoA synthetase. Journal of Molecular Biology, 285(4), 1633–1653. 10.1006/jmbi.1998.2324

12. Weiße, R. H.-J., Faust, A., Schmidt, M., Schönheit, P., & Scheidig, A. J. (2016). Structure of NDP-forming Acetyl-CoA synthetase ACD1 reveals a large rearrangement for phosphoryl transfer. Proceedings of the National Academy of Sciences, 113(5), E519–E528. 10.1073/pnas.1518614113

13. Atalay, N., Tosun, B., Usluer, O., Öztürk, L., Johnson, J., & DeMirci, H. (2023). Cryogenic X-ray crystallographic studies of biomacromolecules at Turkish Light Source “Turkish DeLight.” Turkish Journal of Biology, 47(1), 1–13. 10.55730/1300-0152.2634

14. Kabsch, W. (2010). XDS. Acta Crystallographica Section D: Biological Crystallography, 66(2), 125–132. 10.1107/S0907444909047337

15. Tickle, I. J., Flensburg, C., Keller, P., Paciorek, W., Sharff, A., Vonrhein, C., & Bricogne, G. (2016). STARANISO. Global Phasing Ltd. http://staraniso.globalphasing.org/cgi-bin/staraniso.cgi

16. Adams, P. D., Afonine, P. V., Bunkóczi, G., Chen, V. B., Davis, I. W., Echols, N., Headd, J. J., Hung, L.-W., Kapral, G. J., Grosse-Kunstleve, R. W., McCoy, A. J., Moriarty, N. W., Oeffner, R., Read, R. J., Richardson, D. C., Richardson, J. S., Terwilliger, T. C., & Zwart, P. H. (2010). PHENIX: A comprehensive Python-based system for macromolecular structure solution. Acta Crystallographica Section D: Biological Crystallography, 66(2), 213–221. 10.1107/s0907444909052925

17. Emsley, P., & Cowtan, K. (2004). Coot: Model-building tools for molecular graphics. Acta Crystallographica Section D: Biological Crystallography, 60(12), 2126–2132. 10.1107/s0907444904019158

18. Schrödinger, LLC. (2021). The PyMOL Molecular Graphics System (Version 2.5.2) [Computer software]. https://pymol.org

19. Altschul, S. F., Madden, T. L., Schäffer, A. A., Zhang, J., Zhang, Z., Miller, W., & Lipman, D. J. (1997). Gapped BLAST and PSI-BLAST: A new generation of protein database search programs. Nucleic Acids Research, 25(17), 3389–3402. 10.1093/nar/25.17.3389

20. Altschul, S. F., Gish, W., Miller, W., Myers, E. W., & Lipman, D. J. (1990). Basic local alignment search tool. Journal of Molecular Biology, 215(3), 403–410. 10.1016/S0022-2836(05)80360-2

21. Tamura, K., Stecher, G., & Kumar, S. (2021). MEGA11: Molecular Evolutionary Genetics Analysis version 11. Molecular Biology and Evolution, 38(7), 3022–3027. 10.1093/molbev/msab120

22. Tamura, K., Peterson, D., Peterson, N., Stecher, G., Nei, M., & Kumar, S. (2011). MEGA5: Molecular evolutionary genetics analysis using maximum likelihood, evolutionary distance, and maximum parsimony methods. Molecular Biology and Evolution, 28(10), 2731–2739. 10.1093/molbev/msr121

23. Letunic, I., & Bork, P. (2021). Interactive Tree Of Life (iTOL) v5: An online tool for phylogenetic tree display and annotation. Nucleic Acids Research, 49(W1), W293–W296. 10.1093/nar/gkab301

24. Yariv, B., Tzemach, A., Yacoby, I., Chor, B., & Pupko, T. (2023). Using evolutionary data to make sense of macromolecules with a “face-lifted” ConSurf. Protein Science, 32(4), e4582. 10.1002/pro.4582

25. Negi, S. S., Schein, C. H., Oezguen, N., Power, T. D., & Braun, W. (2007). InterProSurf: A web server for predicting interacting sites on protein surfaces. Bioinformatics, 23(24), 3397–3399. 10.1093/bioinformatics/btm513

26. Saier, M. H., Jr., & Reizer, J. (1990). Domain shuffling during evolution of the proteins of the bacterial phosphotransferase system. Research in Microbiology, 141(9), 1033–1038. 10.1016/0923-2508(90)90077-4

27. Oliynyk, M., Brown, M. J. B., Cortés, J., Staunton, J., & Leadlay, P. F. (1996). A hybrid modular polyketide synthase obtained by domain swapping. Chemistry & Biology, 3(10), 833–839. 10.1016/s1074-5521(96)90069-1

28. Levy, E. D., & Teichmann, S. A. (2013). Structural, evolutionary, and assembly principles of protein oligomerization. In P. M. Conn (Ed.), Progress in molecular biology and translational science (Vol. 117, pp. 25–51). Academic Press. 10.1016/b978-0-12-386931-9.00002-7

29. Ren, M., Feng, X., Huang, Y., Wang, H., Hu, Z., Clingenpeel, S., Swan, B. K., Ferriera, S., Giglio, S., Li, G., Wu, L., Yan, J., Yang, Y., Zhang, T., Stepanauskas, R., Woyke, T., & Luo, H. (2019). Phylogenomics suggests oxygen availability as a driving force in Thaumarchaeota evolution. The ISME Journal, 13(9), 2150–2161. 10.1038/s41396-019-0415-1

30. Verschueren, K. H., Blanchet, C., Felix, J., Dansercoer, A., De Vos, D., Bloch, Y., Van Beeumen, J., Svergun, D. I., Gutsche, I., & Verstraete, K. (2019). Structure of ATP citrate lyase and the origin of citrate synthase in the Krebs cycle. Nature, 568(7753), 571–575. 10.1038/s41586-019-1095-5

31. Huang, J., & Fraser, M. E. (2016). Structural basis for the binding of succinate to succinyl-CoA synthetase. Acta Crystallographica Section D: Structural Biology and Crystallization Communications, 72(8), 912–921. 10.1107/S205979831601131X

32. Fraser, M. E., Joyce, M. A., Ryan, D. G., & Wolodko, W. T. (2002). Two glutamate residues, Glu 208 alpha and Glu 197 beta, are crucial for phosphorylation and dephosphorylation of the active-site histidine residue in succinyl-CoA synthetase. Biochemistry, 41(2), 537–546. 10.1021/bi011700g

33. Joyce, M. A., Fraser, M. E., James, M. N., Bridger, W. A., & Wolodko, W. T. (2000). ADP-binding site of *Escherichia coli* succinyl-CoA synthetase revealed by x-ray crystallography. Biochemistry, 39(1), 17–25. 10.1021/bi991901a

34. Bailey, D. L., Fraser, M. E., Bridger, W. A., James, M. N., & Wolodko, W. T. (1999). A dimeric form of *Escherichia coli* succinyl-CoA synthetase produced by site-directed mutagenesis. Journal of Molecular Biology, 285(4), 1655–1666. 10.1006/jmbi.1998.2325

35. Fan, F., Liu, Y., Zhao, F., & Wang, Q. A. (2012). On the catalytic mechanism of human ATP citrate lyase. Biochemistry, 51(26), 5198–5211. 10.1021/bi3003713

